# Proximity-dependent biotinylation and identification of flagellar proteins in *Trypanosoma cruzi*

**DOI:** 10.1101/2023.02.16.528900

**Authors:** Madalyn M. Won, Aaron Baublis, Barbara A. Burleigh

## Abstract

The flagellated kinetoplastid protozoan and causative agent of human Chagas disease, *Trypanosoma cruzi*, inhabits both invertebrate and mammalian hosts over the course of its complex life cycle. In these disparate environments, *T. cruzi* uses its single flagellum to propel motile life stages and in some instances, to establish intimate contact with the host. Beyond its role in motility, the functional capabilities of the *T. cruzi* flagellum have not been defined. Moreover, the lack of proteomic information for this organelle, in any parasite life stage, has limited functional investigation. In this study, we employed a proximity-dependent biotinylation approach based on the differential targeting of the biotin ligase, TurboID, to the flagellum or cytosol in replicative stages of *T. cruzi*, to identify flagellar-enriched proteins by mass spectrometry. Proteomic analysis of the resulting biotinylated protein fractions yielded 218 candidate flagellar proteins in *T. cruzi* epimastigotes (insect stage) and 99 proteins in intracellular amastigotes (mammalian stage). Forty of these flagellar-enriched proteins were common to both parasite life stages and included orthologs of known flagellar proteins in other trypanosomatid species, proteins specific to the *T. cruzi* lineage and hypothetical proteins. With the validation of flagellar localization for several of the identified candidates, our results demonstrate that TurboID-based proximity proteomics is an effective tool for probing subcellular compartments in *T. cruzi.* The proteomic datasets generated in this work offer a valuable resource to facilitate functional investigation of the understudied *T. cruzi* flagellum.

**Importance:** *Trypanosoma cruzi* is a protozoan parasite that causes Chagas disease, which contributes substantial morbidity and mortality in South and Central America. Throughout its life cycle, *T. cruzi* interacts with insect and mammalian hosts via its single flagellum, establishing intimate contact with host membranes. Currently, few flagellar proteins have been identified in *T. cruzi* that could provide insight into the mechanisms involved in mediating physical and biochemical interactions with the host. Here, we set out to identify flagellar proteins in the main replicative stages of *T. cruzi* using a proximity-labeling approach coupled with mass spectrometry. The >200 candidate flagellar proteins identified represent the first large scale identification of candidate flagellar proteins in *T. cruzi* with preliminary validation. These data offer new avenues to investigate the biology of *T. cruzi* - host interactions, a promising area for development of new strategies aimed at the control of this pathogen.

## Introduction

*Trypanosoma cruzi* is the uniflagellate protozoan parasite that causes Chagas disease, a chronic disease with severe outcomes including cardiomyopathies and gastrointestinal motility disorders^1,2^. *T. cruzi* has a complex life cycle that involves both invertebrate and mammalian hosts, in which the parasite undergoes marked developmental changes and alternates between actively dividing (‘epimastigote’ or ‘amastigote’ forms in insect and mammalian hosts, respectively) and non-dividing ‘trypomastigote’ forms in both hosts (life cycle schematic; **Supplementary Fig. 1**). In mammals, infection is initiated by motile trypomastigotes that actively invade host cells before converting to the non-motile amastigote stage that replicates in the host cytoplasm. Intracellular *T. cruzi* amastigotes begin to replicate ~24 hours post-infection (hpi) and undergo several rounds of cell division before converting back to trypomastigotes that eventually rupture the host cell membrane (between ~90-120 hpi) to allow dissemination of the parasite and infection of new tissue sites. Once *T. cruzi* infection is established in mammalian hosts, parasites typically persist at low levels for the life of the host, giving rise to chronic infections that can trigger inflammation and pathology.

In both insect and mammalian hosts, *T. cruzi* can establish intimate contact with host structures using its single flagellum ^3–5^. In triatomine vectors, epimastigotes attach to the hindgut by forming a hemidesmosome-like structure between the distal part of the flagellum and host rectal epithelium ^5^. This attachment prevents the parasites from being flushed from the insect and is important for promoting differentiation to the infectious metacyclic stage ^5^. In mammalian host cells, cytosolically-localized *T. cruzi* amastigotes establish intermittent contact with host mitochondria using their short motile flagellum ^3,6^. Unlike the motile trypomastigote and epimastigote stages of *T. cruzi*, that have elongated flagella (up to 15 μm in length ^7^), replicative intracellular amastigotes have a truncated flagellum (~2.7 μm) that extends just beyond the opening of the flagellar pocket ^6^. Also, *T. cruzi* amastigotes retain a 9+2 axonemal structure found in motile trypanosomatid life stages ^8^, but lack a paraflagellar rod, a unique lattice-like structure that runs parallel to the axoneme in these organisms ^9^, and which is associated with several functions including flagellar motility and signal transduction ^10^. It has long been assumed that the minimal amastigote flagellum serves no function other than to provide a structural platform for flagellar outgrowth during differentiation to motile life stages ^11^. However, recent observations that the flagellum of intracellular *T. cruzi* amastigotes undergoes low frequency aperiodic ‘beating’ inside mammalian host cells ^6^ and makes physical contact with the host mitochondria ^3,6^, indicate that the amastigote flagellum has a functional role within the host cell. The interaction between the *T. cruzi* amastigote flagellum and host mitochondria is comparable to the intimate contact observed between the flagellum of intracellular *Leishmania mexicana* amastigotes and the host parasitophorous vacuole membrane ^12^. In the case of *Leishmania*, it has been postulated that the amastigote flagellum has a sensory role and/or functions in the delivery of parasite material to the infected host cell ^12,13^. It is therefore reasonable to predict that the *T. cruzi* amastigote flagellum may have similar role(s) in its interactions with the intracellular environment of the host cell.

In addition to critical roles in motility, eukaryotic flagella (i.e. cilia) and non-motile cilia have emerged as important sensory organelles that are equipped with signal transduction systems and second messengers such as cyclic AMP (cAMP) ^14^ and calcium ^15,16^ that coordinate cellular responses to external stimuli. Functions beyond cell locomotion have also been ascribed to the flagellum of motile trypanosomatid life stages ^11,13,17–19^, where the best understood example of sensory integration in these organisms is the role of flagellar receptor-type adenylate cyclases and cAMP-depending signaling in pH taxis and social motility in the insect stage of *Trypanosoma brucei* ^20–23^. In *Leishmania*, flagellar aquaporin has been implicated in osmotaxis ^24^ in the insect stage, and the flagellar membrane may be a critical site for glucose ^25,26^ and arginine ^27^ sensing in these parasites. Indeed, in both *T. brucei* and *Leishmania*, near comprehensive flagellar proteomes have been generated using shot-gun proteomics of isolated flagella ^28–30^ or of detergent and high salt extracted fractions of the parasite, yielding axonemal and paraflagellar rod proteins^31^. Further, in *T. brucei*, specific domains of the flagellum have been partially mapped using proximity-dependent biotinylation including flagellar attachment zone proteins ^32^ and the flagellar tip ^33^, a specialized signaling domain.

By comparison, we have little knowledge of the molecular composition of the *T. cruzi* flagellum. Beyond a core axonemal proteome that is predicted based on conservation across trypanosomatid species and life stages ^34^, few flagellar proteins that have the potential to serve as a functional interface with the host environment have been identified in any *T. cruzi* life stage ^9,35,36^. The best characterized is the flagellar calcium-binding protein (FCaBP), a dual-acylated, 24 kDa Ca^2+^-sensing protein that tethers to the inner leaflet of the flagellar membrane ^37^. FCaBP is expressed in all *T. cruzi* life stages and is conserved across other trypanosomatid species, but its precise role in the biology of these organisms is unknown beyond its role as a calcium binding protein ^36,38,39^. Additionally, we have recently localized small myristoylated protein 1-1 (TcSMP1-1) to the flagellum in amastigotes ^6^, but the overall proteomic landscape of the *T. cruzi* flagellum remains largely uncharacterized.

In this study, we pursued a targeted, proximity-dependent biotinylation (BioID) approach to identify flagellar membrane and membrane-proximal flagellar proteins in the replicative stages of *T. cruzi.* We report the identification of 218 and 99 candidate flagellar proteins in *T. cruzi* epimastigotes and intracellular amastigote stages, respectively, many of which are conserved in other trypanosomatid species with evidence of flagellar localization. Approximately 20% of the candidate flagellar proteins were found to be restricted to the *T. cruzi* lineage, including a hypothetical protein that we confirmed localizes to the flagellar tip in *T. cruzi* epimastigotes and intracellular amastigotes. The novel BioID dataset identified here provides a critical foundation for investigation of the *T. cruzi* flagellum and its role in mediating interactions with diverse host environments.

## Results

### Flagellar and cytosolically targeted TurboID retains activity in *T. cruzi*

To facilitate the identification of flagellar proteins in *T. cruzi* using a proximity-dependent biotinylation approach, we generated transgenic parasites that express the biotin ligase, TurboID ^40^ in the parasite flagellum, as an in-frame fusion with C-terminal FLAG-tagged *T. cruzi* small myristoylated protein 1-1 (TcSMP1-1) (**Fig. 1A,B;** SMP1-1-FLAG-TurboID; ‘*F-Turbo*’). TcSMP1-1 was chosen as the endogenous ‘bait’ protein for flagellar localization of TurboID given its near exclusive localization in the flagellum in both replicative stages of *T. cruzi*, epimastigotes and amastigotes ^6^ (**Fig. 1A**), and because TcSMP1-1 contains the N-myristoylation sequence motif (MGXXXS/T) required for localization and tethering to the inner flagellar membrane, as demonstrated in *Leishmania* ^41^. The strategy of targeting TurboID to the flagellum using TcSMP1-1 is expected to increase the likelihood of identifying flagellar membrane and associated proteins while minimizing capture of axonemal proteins. To control for non-flagellar TurboID expression in the F-Turbo parasites, we generated an independent transgenic line that expresses FLAG-TurboID in the cytoplasm (**Fig. 1B**; ‘*C-Turbo*’). The parallel processing of F-Turbo and C-Turbo parasites, along with parental (WT) parasites that lack TurboID expression (**Fig. 1C)**, will allow for background subtraction and identification of flagellar-enriched proteins in F-Turbo versus C-Turbo within the same parasite life cycle stage.

**Figure 1.**
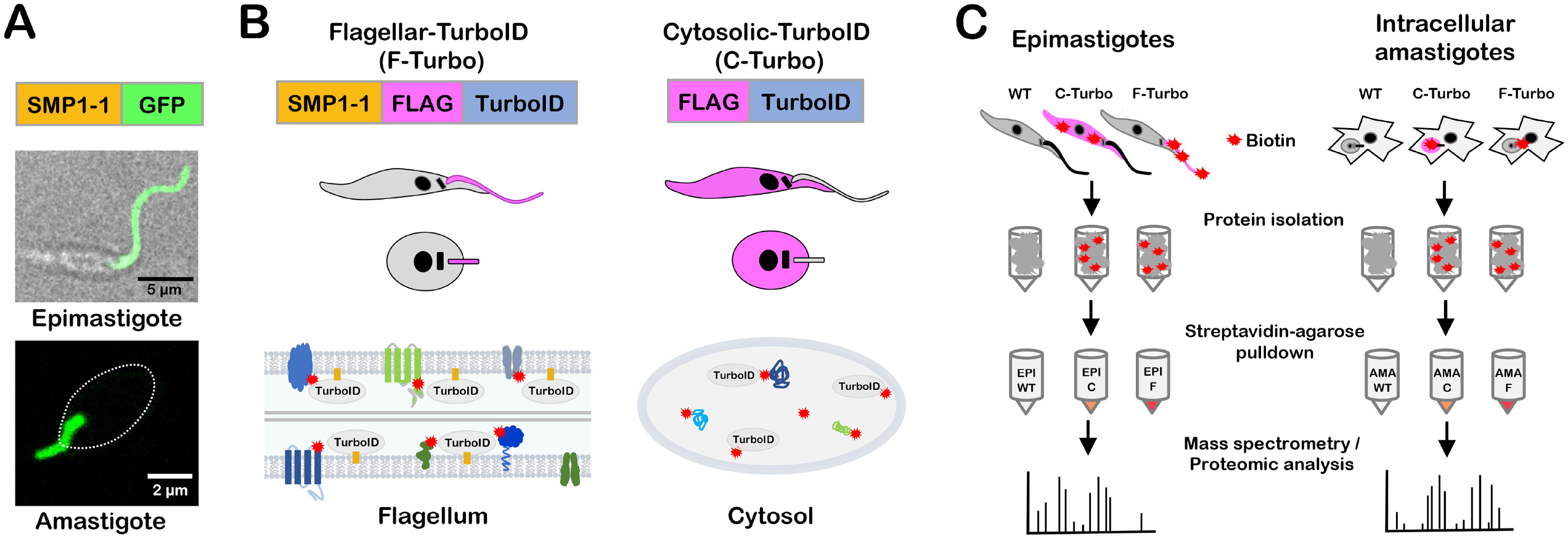
*T. cruzi* life cycle and schematic of TurboID-expressing lines generated for proximity-dependent biotinylation experiments. (**A**) Live confocal images of SMP1-1-GFP localized to the flagellum of *T. cruzi* epimastigotes and an intracellular amastigote; white oval denotes position of amastigote body. (**B**) Strategy for generating stable *T. cruzi* lines expressing TurboID in the flagellum using SMP1-1 as the endogenous bait protein or in the cytoplasm of epimastigotes and amastigotes, where addition of exogenous biotin will mediate biotinylation (red star) of proteins in close proximity to TurboID in both settings. FLAG-epitope is included to facilitate TurboID localization in transfectants. (**C**) Flow chart outlining the experimental protocol used for identification of biotinylated proteins in epimastigotes (*left*) and intracellular amastigotes (*right*). For both life stages, wild-type (*‘WT’*), cytoplasmic-TurboID (‘*C’*) and flagellar-TurboID (‘*F’*) parasites (from left to right in the illustration) were exposed to biotin and the biotinylated protein fraction in protein lysates captured on streptavidin-agarose beads and subjected to mass spectrometry for identification and subsequent proteomic analysis.

TurboID expression in transgenic *T. cruzi* parasites was confirmed by indirect immunofluorescence microscopy of fixed parasites stained with an antibody to the FLAG tag epitope, located immediately upstream of TurboID (**Fig. 2**). The flagella of F-Turbo parasites were brightly stained (**Fig. 2A,C;** *F-Turbo*) indicating that trafficking of TurboID to the flagellum occurred in both *T. cruzi* epimastigote (**Fig. 2A;** *F-Turbo*) and amastigote (**Fig. 2C;** *F-Turbo*) life stages. While most of the FLAG signal was localized to the flagellum in epimastigotes (**Fig. 2A;** *F-Turbo*), signal was detected in the body of intracellular amastigotes in addition to the brightly stained flagellum (**Fig. 2C;** *F-Turbo*), which may be due to overexpression of the SMP1-1-FLAG-TurboID fusion protein. C-Turbo parasites, generated as a proteomic control for non-flagellar TurboID-dependent biotinylation (**Fig. 1C**), were confirmed to express cytosolic FLAG in both parasite life stages (**Fig. 2A,C;** *C-Turbo*). To determine if TurboID is active in *T. cruzi*, total protein lysates were prepared from WT and Turbo-expressing parasites, following brief exposure to exogenous biotin, were probed with streptavidin-DyLight™ 800 to detect biotinylated proteins (**Fig. 2B,D**). As expected, multiple biotinylated proteins were revealed in lysates derived from F-TurboID and C-TurboID epimastigotes (**Fig. 2B**) and amastigotes (**Fig. 2D**) whereas few biotinylated proteins were detected in the parental (WT) controls. Differences in the biotinylated protein profiles observed when comparing F-Turbo to C-Turbo within a single life stage (**Fig. 2B,D)** likely reflect the differential localization patterns for TurboID in these parasite lines. Combined, these results confirm the expression of active TurboID in the flagellum (F-Turbo) or cytosol (C-Turbo) in both replicative stages of *T. cruzi.*

**Figure 2.**
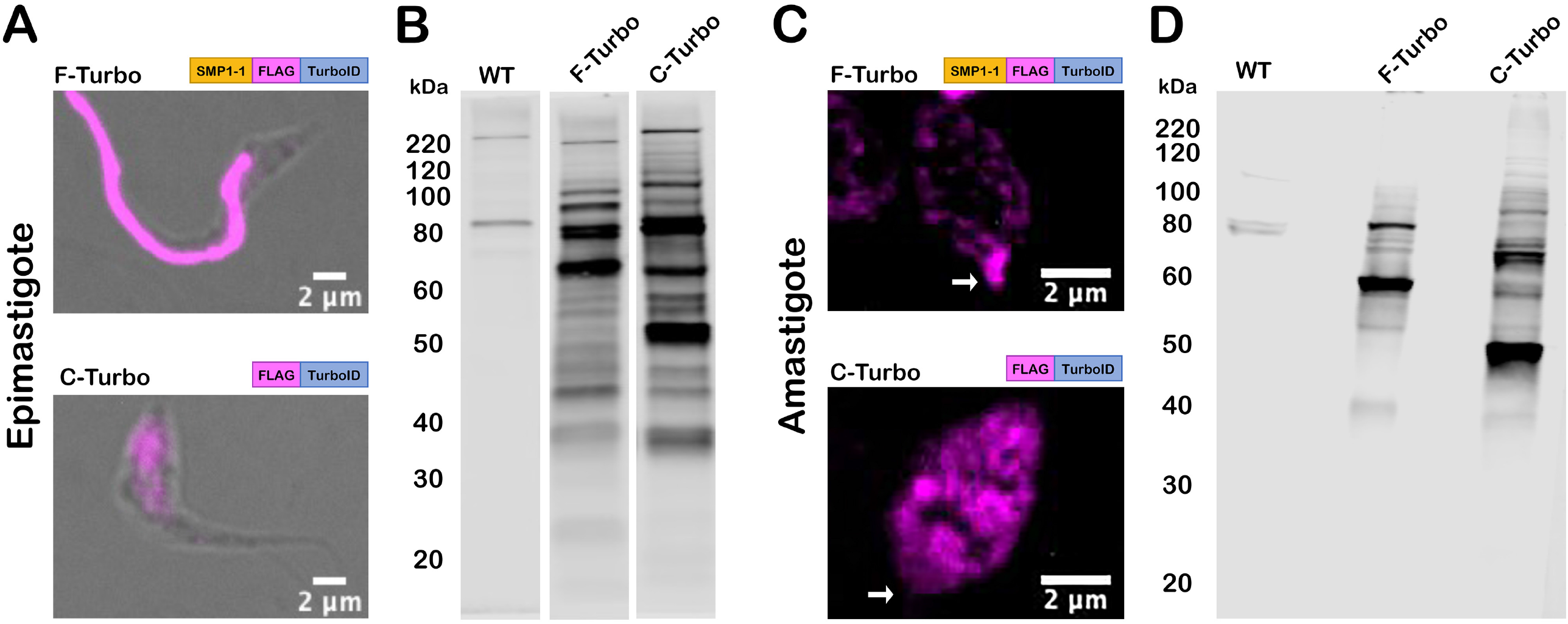
TurboID localization and activity in *T. cruzi*. (**A,C**) Fluorescence microscopy images of fixed *T. cruzi* epimastigotes (**A**) or amastigotes (**C**) expressing SMP1-1-FLAG-TurboID (F-Turbo) (*top*) or FLAG-TurboID (C-Turbo) (*bottom*) stained for FLAG epitope (anti-FLAG)(*pink*). In (**C**), white arrows indicate the position of the amastigote flagellum. (**B,D**) Biotinylated proteins in lysates of WT, F-Turbo, and C-Turbo *T. cruzi* epimastigote (**B**) and amastigotes (**D**) detected with streptavidin-Dylight800.

### Proteomic identification of candidate flagellar proteins in *T. cruzi*

Biotinylated proteins in lysates generated from WT, F-Turbo and C-Turbo *T. cruzi* epimastigotes or intracellular amastigotes, were captured on immobilized streptavidin beads and identified using high performance liquid chromatography combined with mass spectrometry (**Fig. 1C**). Three independent biological replicates were analyzed for each parasite line, with the exception of F-Turbo epimastigotes, for which triplicate samples from two independent transfections were included. Peptide identification and relative intensity data obtained for replicate samples from each parasite line are represented in **Supplementary Table 1**. Principal component analysis (PCA) identified overall trends in the proteomic data obtained for *T. cruzi* epimastigotes (**Fig. 3A**) and amastigotes (**Fig. 3C**), revealing that biological replicates from individual parasite lines (WT, F-Turbo, C-Turbo) formed discrete clusters that were well separated from each other. As the replicates from independent F-Turbo epimastigote lineages were indistinguishable, these samples were pooled for subsequent analyses. Prior to data filtering and analysis, protein intensity scores were averaged across biological replicates within individual experimental groups (**Supplementary Table 1**).

**Figure 3.**
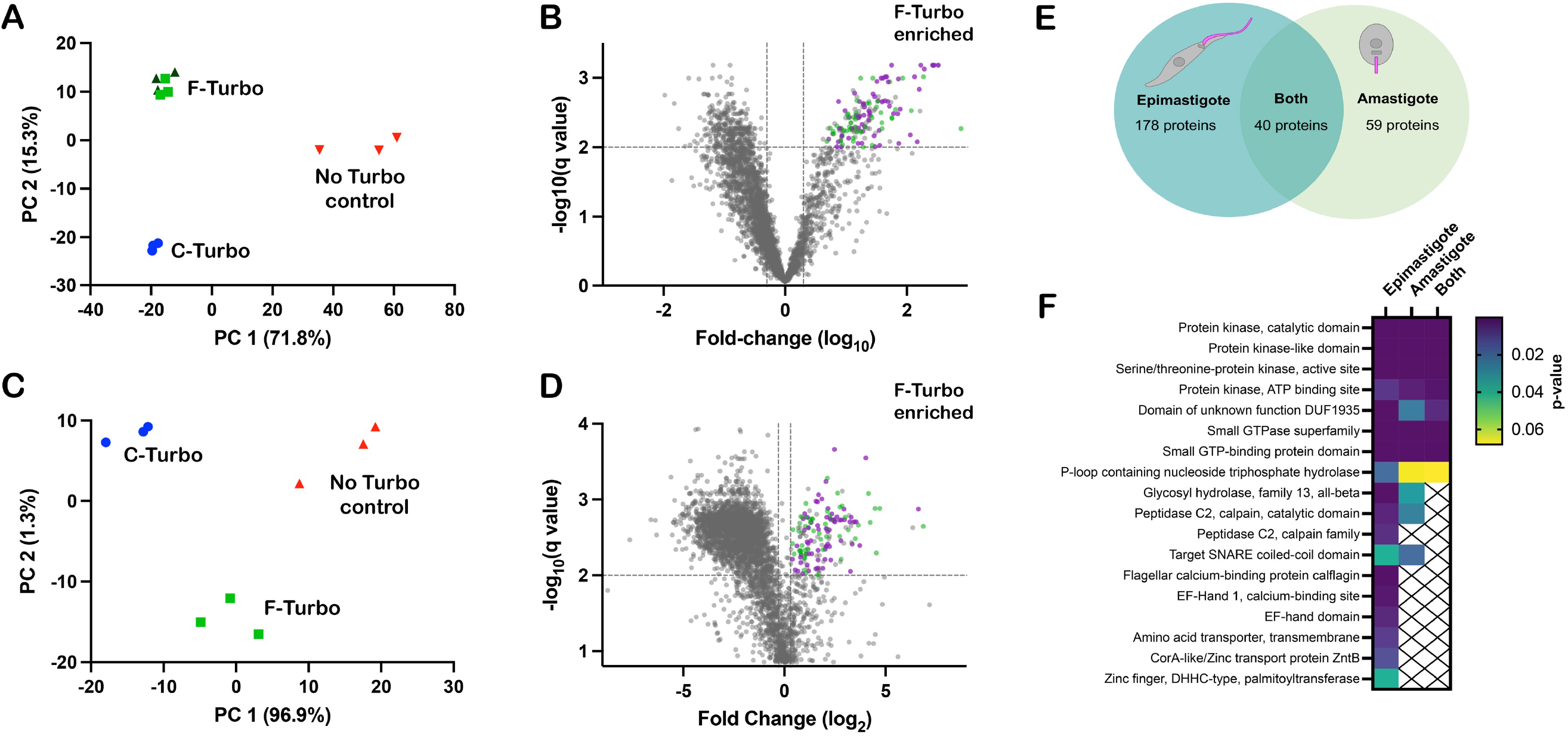
Proximity proteome analysis identifies flagellar-enriched proteins in *T. cruzi*. Principal component analysis (PCA) of biotinylome data plotted for WT (*no TurboID control*), flagellar-TurboID (*F-Turbo*) and cytosolic-TurboID (*C-Turbo*) for *T. cruzi* epimastigotes (**A**) and intracellular amastigotes (**C**). The two independent F-Turbo groups in the epimastigote PCA plot are represented in green (triangles and squares). Volcano plots (**B,D**) with fold-change (F-Turbo vs C-Turbo; x-axis) and adjusted p-value (q-value; y-axis) for *T. cruzi* epimastigote (**B**) and amastigote (**D**) proteomic data. Horizontal lines represent a q-value of 0.01 and the two vertical lines indicate the cut-offs for fold change (2-fold). The top right quadrants in each plot (**B,D**) contain proteins that are significantly enriched in F-Turbo proteomes (q<0.01, >2-fold change). Known trypanosomatid flagellar proteins (*purple circles*) and hypothetical proteins (*green circles*) are shown for the F-Turbo enriched proteins. (**E**) Venn diagram depicting the number of proteins identified as enriched in the F-Turbo samples of *T. cruzi* epimastigote and amastigote stages. **(F**) Interpro domains assigned by DAVID that are significantly enriched in F-Turbo samples (i.e. proteins found in the upper right quadrant of each volcano plot) in epimastigotes, amastigotes and those common to both life stages; *p*-value is a modified Fisher exact, for protein enrichment analysis.

Streptavidin-bound proteins identified by mass spectrometry in WT parasites (which lack TurboID) represent the ‘background’ signal of endogenously biotinylated proteins and proteins that bound non-specifically to immobilized streptavidin. Thus, proteins represented at less than 100-fold enriched over the WT samples in F-Turbo or C-Turbo epimastigote samples were removed before subsequent analysis. Also, any proteins identified in less than 4/6 of the F-Turbo samples were removed (**Supplementary Table 2**). Volcano plots revealed the protein subsets significantly enriched in F-Turbo and C-Turbo samples in epimastigotes (**Fig. 3B**) and amastigotes (**Fig. 3D**). Proteins found to be significantly enriched in F-Turbo over C-Turbo (fold-change > 2; q-value ≤ 0.01) as well as proteins identified uniquely in F-Turbo samples (i.e., not present in C-Turbo samples from the same parasite life stage) are listed in **Supplementary Table 3**. From this analysis, 218 proteins were identified as significantly enriched in F-Turbo samples from *T. cruzi* epimastigotes and 99 proteins in amastigotes (**Supplementary Table 3**).

### The *T. cruzi* SMP1-1 proximity proteome includes known trypanosomatid flagellar proteins

The searchable TrypTag database ^42^, which contains localization data for 7,487 *T. brucei* proteins, was used as a resource to identify orthologues in the *T. cruzi* flagellar-enriched protein dataset (**Supplementary Table 3**) that have demonstrated flagellar localization in *T. brucei.* Of the 218 flagellar-enriched proteins in *T. cruzi* epimastigotes, 145 have orthologs that are represented in the TrypTag database and of these, 75 proteins exhibit at least partial flagellar localization in *T. brucei* bloodstream forms ^42^. Similar results emerged from the *T. cruzi* amastigote data where orthologs of 75 of the 99 proteins found to be enriched in amastigote F-Turbo samples had orthologs in *T. brucei* and were endogenously tagged, 44 of these showed at least partial flagellar localization. A comparison of the flagellar-enriched proteins identified in both *T. cruzi* epimastigotes and amastigotes revealed 40 proteins common to both life stages (**Fig. 3E**), of which 29 have orthologs that are represented in the TrypTag database (**Supplementary Table 3**) and 20 proteins exhibited some flagellar localization in *T. brucei.* Examples of confirmed flagellar proteins in other trypanosomatids that are significantly enriched in the *T. cruzi* flagellar proximity proteome include: flagellar membrane 8 ^43^, flabarin ^44^, flagellar attachment zone 14 ^32^, casein kinase I ^42^, CARP3 ^20^ and cysteine peptidase, Clan CA, family C2 (calpain 1.3) ^42^. Although a significant proportion of the flagellar candidates identified in *T. cruzi* epimastigotes and amastigotes fall into the ‘hypothetical’ category (i.e., no annotation), the datasets were found to be enriched in kinase domains, calpain domains, and small GTP binding protein / GTPase domains (**Fig. 3F**).

### Selected flagellar candidates localize to the *T. cruzi* flagellum in epimastigotes and amastigotes

To localize candidate flagellar proteins in *T. cruzi*, we prioritized those that were significantly enriched in both the epimastigote and amastigote F-Turbo datasets and that included one or more of the following characteristics: (a) sequence motifs predicting membrane localization; (b) predicted role in signaling based on annotation or (c) were unique to the *T. cruzi* lineage (i.e., no obvious orthologues in other trypanosomatid species). Based on these criteria, six proteins were selected for endogenous FLAG-tagging and subsequent subcellular localization (**Table 1**) using primers and homology-directed repair templates shown in (**Supplementary Fig. 4**). Four of the 6 proteins were successfully tagged and three of which exhibited flagellar localization in *T. cruzi* epimastigotes: calpain 1.3 (TcCLB.506563.200), CARP3 (TcCLB.506681.40) and hypothetical protein (TcCLB.510329.180) (**Fig. 4A; Supplementary Fig. 2)**. Another hypothetical protein (TcCLB.509965.20) was not verified as flagellar as the FLAG epitope signal localized to the parasite body (data not shown). Calpain 1.3-mRuby2-smFP FLAG exhibited a punctate pattern of labeling along the entire length of the *T. cruzi* epimastigote flagellum (**Fig. 4A**), whereas expression appeared to be restricted to the flagellar tip in the intracellular amastigote stage (**Fig. 4B**). Both CARP3 and the hypothetical protein (TcCLB.510329.180) localized to the distal region of the epimastigote flagellum (**Fig. 4A**) and hypothetical protein (TcCLB.510329.180) also localized to the flagellar tip in amastigotes (**Fig. 4B**). We were unable to determine CARP3 localization in amastigotes, due to undetectable signal for CARP3-mRuby2-smFP FLAG expression in this life stage, despite clear signal and flagellar localization in epimastigotes (**Fig. 4A**). Nonetheless, the successful identification of a subset of flagellar-localized proteins in intracellular *T. cruzi* amastigotes and epimastigotes provides initial validation of the proteomic datasets generated from proximity-dependent labeling.

**Figure 4.**
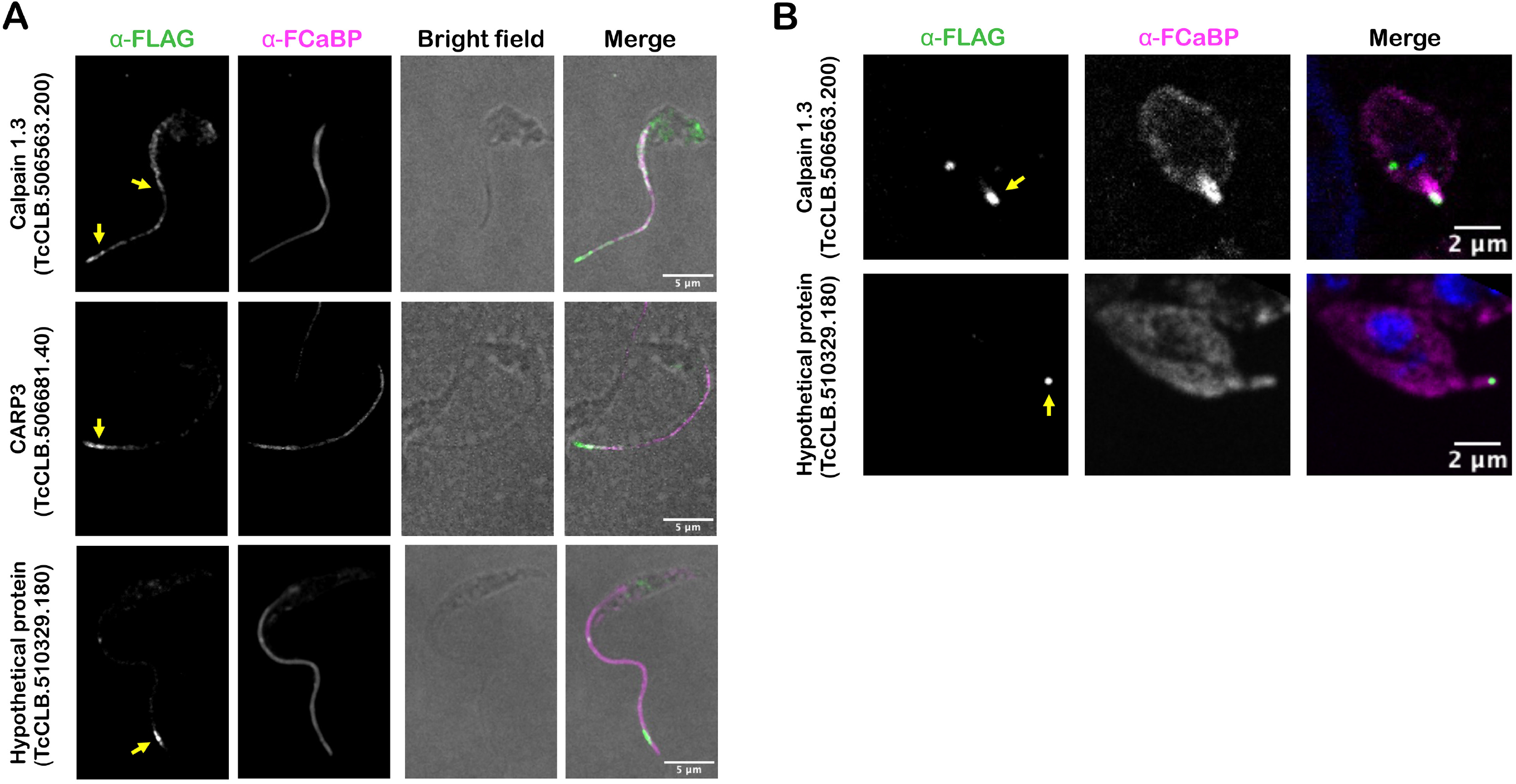
Flagellar localization of candidate flagellar proteins in *T. cruzi*. Endogenous tagging reveals flagellar localization of candidate flagellar proteins: calpain 1.3-smFLAG (TcCLB.506563.200), CARP3-smFLAG (TcCLB.506681.40), or hypothetical protein-FLAG (TcCLB.510329.180) in *T. cruzi* epimastigotes (**A**) or intracellular amastigotes (**B**). In all cases, the FLAG tag was detected in fixed parasites using an anti-FLAG antibody and secondary antibody (*green*) and the flagellum was detected using anti-FCaBP and secondary antibody (*magenta*). The FLAG signal in the flagellum is indicated (*yellow arrow*).

**Table 1:** Candidate flagellar proteins in *T. cruzi* selected for endogenous tagging.

| Name | TriTryp ID | Successfully Tagged? | Reason for selection |
| --- | --- | --- | --- |
| Calpain 1.3 | TcCLB.506563.200 | Yes | Potential signaling role |
| CARP3 | TcCLB.506681.40 | Yes | Potential signaling role |
| Hypothetical Protein | TcCLB.510329.180 | Yes | No known ortholog |
| Hypothetical Protein | TcCLB.509965.20 | Yes | No known *T. brucei* ortholog |
| ATPase | TcCLB.506925.410 | No | Potential transmembrane protein |
| Hypothetical Protein | TcCLB.509011.50 | No | Potential signaling role |

Candidates from the 40 proteins identified in both *T. cruzi* epimastigotes and amastigotes selected for endogenous epitope tagging and localization.

## Discussion

In the present work we demonstrate the successful use of a proximity labeling tool in the protozoan parasite, *Trypanosoma cruzi.* With the goal of identifying flagellar membrane and/or associated proteins in *T. cruzi* as candidates for mediating physical or functional interactions with insect or vertebrate hosts, the biotin ligase TurboID was targeted to the parasite flagellum as a fusion protein with the inner flagellar membrane protein, SMP1-1. Overexpression of the fusion protein was well-tolerated in the parasite, with no interference in the ability to transition between axenic epimastigotes, the *T. cruzi* life stage in which DNA transfection and selection is performed, and the intracellular stages in mammalian cells. This offered the opportunity to perform a comparative analysis of the two main replicative stages of *T. cruzi*, one that is motile with an elongated flagellum (epimastigote) and the other that is non-motile with a short flagellum (intracellular amastigote). Furthermore, inclusion of cytosolic-TurboID expressing parasites in the analysis aided in the differential identification of flagellar-enriched biotinylated proteins derived from SMP1-1-FLAG-TurboID parasites. This approach yielded 218 flagellar candidates in epimastigotes and 99 proteins in amastigotes, where 40 proteins were common to both *T. cruzi* life stages. Flagellar localization was confirmed for a subset of proteins in this dataset, based on endogenous epitope-tagging, and many more were predicted based on demonstrated localization in the related trypanosomatids, *T. brucei* or *Leishmania spp.*

The functional capabilities of the *T. cruzi* flagellum are broadly uncharacterized, beyond its role in propelling motile life stages and anchoring epimastigotes to the rectal mucosa in the insect vector ^4,45,46^. However, the recent recognition that the flagellum of cytosolically-localized intracellular amastigotes is capable of beating and establishes physical contact with host mitochondria ^3,6^, points to a potential role for the amastigote flagellum in host environmental sensing. Although little is known regarding the sensory capabilities of trypanosomatids in general, significant progress has been made toward a molecular understanding of pH taxis and social motility in the insect stages of *T. brucei* ^21,23^. The sensory system involves regulation of cyclic AMP levels, modulated by flagellar receptor-type adenylate cyclases and cyclic AMP phosphodiesterases^22^ and the involvement of a cyclic AMP responsive protein (CARP3), which is thought to act in a complex with adenylate cyclases ^20^. Our discovery, that CARP3 is expressed in the replicative stages of *T. cruzi*, including intracellular amastigotes (despite the inability to localize the tagged protein in this life stage), is quite exciting given the established role of CARP3 in *T. brucei* ^20^. In *T. brucei* CARP3 is known to co-localize with calpain 1.3 at the distal region of the flagellum, where the two proteins may physically interact ^20^. We show that the calpain 1.3 ortholog is also expressed in the flagellum of *T. cruzi*, where it localizes exclusively to the flagellar tip in intracellular amastigotes, a recognized signaling domain in trypanosomatids ^33(p2),47^. Calpain 1.3 belongs to a sub-family of cysteine peptidases that are predicted to be catalytically inactive as they lack one or more of the active site amino acid residues ^48^. Proteins with these features are thought to play a role in calcium homeostasis / signaling, including the calcium-based regulation of adenylate cyclase complexes ^20,49^. Despite the lack of social motility in *T. cruzi*, the expression of flagellar CARP3 and calpain 1.3 in this species is a strong indicator that the *T. cruzi* flagellum is equipped to sense and integrate signals from the environment. Dissection of the functional roles of CARP3 and calpain 1.3 in *T. cruzi* is expected to be instrumental in establishing the existence of flagellum-based environmental sensing in this parasite. In addition, a functional investigation of the hypothetical protein (TcCLB.509965.20) that also localizes to the distal end of the *T. cruzi* flagellum but lacks an obvious ortholog in *T. brucei* or *Leishmania*, has the potential to reveal novel biological or mechanistic insights into the role of the *T. cruzi* flagellum in different life stages.

The proteomic datasets generated here offer new opportunities to pursue the localization and functional analyses of many uncharacterized proteins, some of which are unique to the *T. cruzi* lineage. While some of the annotated proteins are not predicted to localize to the flagellum (based on annotation and TrypTag localization), including proteins involved in protein trafficking which may have encountered SMP1-1-FLAG-TurboID *en route* to the flagellum, there are a number of proteins with transmembrane domains or N-myristoylation consensus sequences that predict membrane-association. SMP-1 contains the N-myristoylation sequence motif (MGXXXS/T) known to direct flagellar localization and tethering to the inner flagellar membrane in *Leishmania*, where it associates tightly with detergent-resistant membranes (lipid rafts) ^41^ and forms homodimers^41^. As such, SMP1-1 may preferentially interact with other membrane proteins associated with lipid rafts in the flagellar membrane. It is notable that a number of proteins that were among the strongest ‘hits’ in the flagellar candidate pool, such as calpain 1.3 and CARP3, also have MGXXXS/T motifs. While this motif is insufficient to direct a protein to the flagellum ^41^, proteins with lipid anchoring motifs or transmembrane domains could be prioritized for future studies of the *T. cruzi* flagellum. Notably, we did not identify adenylate cyclases in our data, even though CARP3 and calpain 1.3 were identified in a proximity-labeling study in *T. brucei* designed to identify flagellar tip proteins that interact with adenylate cyclase 1 ^33^. While the proximity-dependent labeling approach used in this study enabled the discovery of a subset of flagellar proteins in *T. cruzi* amastigotes (where physical isolation of the short amastigote flagellum may not be feasible), it is understood that the resulting proteomes derived for the two parasite life stages are not comprehensive and many *T. cruzi* flagellar membrane proteins remain to be identified. With the discovery of additional flagellar proteins in *T. cruzi*, opportunities will be presented to use one or more of these proteins as alternative bait proteins for proximity labeling with a view to expanding the flagellar proteome in this understudied parasite.

Overall, we have presented the first use of proximity-dependent biotinylation in *T. cruzi* for the identification of more than 200 candidate flagellar proteins across two parasite life stages, thereby creating an important resource for the research community. As more information becomes available for the *T. cruzi* amastigote flagellum, it is expected to provide some context for the biological role of the flagellum in infection and these interactions may be the key for specific targeting of parasite function and viability within the mammalian host. Future investigation focused on identifying the function of potential sensory flagellar candidates in *T. cruzi* epimastigote and amastigote flagella, may aid in the discovery of these currently unknown host-parasite interaction mechanisms.

## Methods and Materials

### Reagents

Compounds were purchased and diluted to stock concentrations: Biotin, 100 mM in DMSO (Sigma Aldrich, St. Louis, Missouri, USA). Phenylmethylsulfonyl fluoride (PMSF), 10 mM in isopropanol (Sigma Aldrich, St. Louis, Missouri, USA). Tosyl-L-lysyl-chloromethane hydrochloride (TLCK), 5 mM in DMSO (Abcam, Cambridge, United Kingdom).

### Mammalian cell culture

Normal Human Neonatal Dermal Fibroblasts (NHDF; Lonza, Basel, Switzerland) and monkey kidney epithelial cells (LLC-MK2; American Type Culture Collection) were maintained in Dulbecco’s modified Eagle medium (DMEM; HyClone, Logan, Utah, USA) supplemented with 10% heat-inactivated FBS (Gibco, Waltham, Massachusetts, USA), 25 mM glucose, 2 mM L-glutamine, and 100 U/mL penicillin-streptomycin (DMEM-10) at 37°C and 5% CO_2_.

### Growth and maintenance of *T. cruzi*

*Trypanosoma cruzi* Tulahuén LacZ clone C4 was obtained from the American Type Culture Collection (ATCC, PRA-330; ATCC, Manassas, Virginia, USA). The epimastigote stage was propagated at 28°C in liver infusion tryptose (LIT) medium (4□gIL NaCl, 0.4□g/L KCl, 8□/L Na2HPO4, 2□g/L dextrose, 3□g/L liver infusion broth, 5□g/L tryptose, with 25□mg/L hemin and 10% heat-inactivated FBS). The mammalian cell infection cycle was initiated with metacyclic trypomastigotes arising within stationary phase epimastigote cultures that were shifted from LIT to DMEM + 2% FBS (DMEM-2) for 5 days at 28°C. Metacyclic-enriched cultures were washed in DMEM-2 and incubated with confluent LLC-MK2 monolayers at 37°C, 5% CO_2_ to allow invasion. Mammalian stage trypomastigotes that emerged from infected LLC-MK2 cells (within 5-10 days) were harvested from culture supernatants and used to infect fresh LLC-MK2 monolayers. This cycle was continued on weekly basis to maintain the mammalian-infective stages of *T. cruzi* in culture. For experimental infections, trypomastigotes collected from LLC-MK2 maintenance cultures were pelleted at 2060 x g for 10 minutes and pellets were incubated at 37°C, 5% CO_2_ for 2-4 hours to allow motile trypomastigotes to swim up into the supernatant. Purified trypomastigotes in the supernatant were collected, washed once in DMEM-2 and utilized to infect sub-confluent monolayers of NHDF as indicated.

### Generation of stable *T. cruzi* transfectants

*T. cruzi* strains expressing TcSMP1-1GFP were previously generated ^6^. A plasmid containing the TurboID sequence was a kind gift of Jeffrey Dvorin (Harvard Medical School). Each TurboID construct was used to replace the GFP-P2A-puro cassette in a modified pTREX plasmid ^50,51^ containing either SMP1-1-GFP or GFP alone. The inserts in the pTREX backbone to generate the F-Turbo plasmids were SMP1-1-TurboID-P2A-puro (F-Turbo-P) or SMP-1-1-TurboID-T2A-puro (F-Turbo-T). The inserts and backbone were assembled using the NEB HiFi DNA assembly kit (New England Biolabs, Ipswich, Massachusetts, USA), resulting in the final plasmids. For the cytosolic control, TurboID-P2A-puro was amplified using PCR and then inserted into the pTREX-GFP backbone, replacing GFP between the SpeI and XmaI cut sites using or through restriction enzyme cloning. To generate TurboID-expressing parasites *T. cruzi* epimastigotes were transfected with 15 μg of the respective DNA. Prior to transfection log-phase *T. cruzi* epimastigotes, were pelleted at 2060 x g for 10 minutes, resuspended in 100 μL of Tb BSF buffer ^52^ (4×10^7^ parasites) and placed into a sterile 2 mm gap cuvette with the appropriate DNA and transfected using an Amaxa Nucleofector II (Lonza, Basel, Switzerland; U-33 program). Parasites were immediately transferred to LIT medium for 24 hours before adding 10 μg/mL puromycin (Invivogen, San Diego, California, USA) or 50 μg/mL blasticidin (Invivogen, San Diego, California, USA) for selection.

CRISPR/Cas9-facilitated epitope tagging of genomic loci in *T. cruzi* was performed as described^53^. Briefly, each gene of interest was PCR-amplified from genomic DNA *(T. cruzi* Tulahuen strain) and PCR products sequenced. Two gRNA binding sites near the 3’ region of each gene of interest were identified using EuPaGDT Editing of the previously modified pTREX-n-Cas9 plasmid ^51^ (Addgene plasmid 68708), performed to exchange the previous gRNA sequence was achieved using a Q5 mutagenesis kit (New England Biolabs, Ipswich, Massachusetts, USA). gRNA sequences were inserted into pTREX-n-Cas9 using primers specific to the gene of interest in **Supplementary Table 4**, such that the previous gRNA sequence was replaced. The template for generating homology-directed repair DNA for gene tagging was constructed by inserting a P2A viral skip peptide in frame with a downstream blasticidin-S deaminase (BSD) or puromycin N-acetyl-transferase (puro) and TOPO cloned into a pCR4 backbone (Thermo Fisher, Waltham, MA, United States of America) ^51^. Homology template was amplified from this template using ultramer pairs (**Supplementary Table 4**) that provided 100 bp of homology for the gene of interest, the FLAG tag and 20 bp of homology to template. Parasite were transfected as above with 25 μg of each gRNA-specific pTREX-n-Cas9 plasmid to the gene of interest and 50 μg of homology repair template. Correct integration of the endogenous tag and drug cassette was established via PCR (**Supplementary Fig. 2**).

### Detection of biotinylated protein fractions in *T. cruzi*

#### Epimastigotes

1.5 x 10^8^ epimastigotes were pelleted at 2060 x g for 10 minutes, resuspended in 1 ml of LIT and incubated with 50 μM biotin for 10 minutes at 37°C. Parasites were washed twice with ice-cold PBS then resuspended in 1 ml cell lysis buffer ^55^ (0.5% Nonidet P-40, 500 mM NaCl, 5 mM EDTA, 1 mM DTT, 50 mM Tris-Base, 0.4% SDS, pH 7.4) with Roche cOmplete™ Protease Inhibitor (Sigma-Aldrich, St. Louis, Missouri, USA), 100 μM PMSF and 10 μM TLCK. Lysates were sonicated using 3 pulses of 30 seconds at 100% amplitude (Q700 sonicator, QSONICA, Newton, Connecticut), with 15 sec breaks between to cool the tubes on ice. Samples were centrifuged at 16,000 x g for 20 minutes at 4°C and the supernatant was collected. Aliquots of clarified lysate (3 x 10^6^ parasite equivalents) were resolved by SDS-PAGE (Mini-PROTEAN®TGX protein gel; Bio-Rad, Hercules, California, USA), transferred to PVDF membrane (Immobilon®-FL, MilliporeSigma, Burlington, Massachusetts, USA) and probed with Streptavidin DyLight™ 800 (Thermo Fisher Scientific, Waltham, Massachusetts, USA) to detect biotinylated proteins.

#### Intracellular amastigotes

At 48 hpi, *T. cruzi*-infected NHDF monolayers in T-150 flasks were exposed to 100 μM biotin for 10 minutes at 37°C, 5% CO_2_. Monolayers were then rinsed three times with cold PBS and incubated with 2 mL of cell lysis buffer (as above). Flasks were agitated manually for 5 minutes then cells were scraped and transferred into a tube containing 0.5 μL of benzonase (Sigma-Aldrich, St. Louis, Missouri, USA). Tubes were placed on a rotative wheel for 15 minutes at room temperature, then sonicated as above. Amastigote loading volumes were normalized via Western blot as follows. Equal volumes of serially diluted protein lysates, generated for WT, F-Turbo and C-Turbo infected NHDF, were resolved by SDS-PAGE (Mini-PROTEAN®TGX gels), transferred to PVDF membrane and probed with a rabbit antibody to trypanosome BiP ^56^, followed by α-Rabbit Alexa Flour 647. A LiCor Odessy® CLx imager was used to measure BiP signal in each sample (Image Studio). Relative BiP densities were used to adjust the volumes of each amastigote lysates (confirmed by independent western blots) prior to loading on streptavidin beads.

### Isolation of biotinylated proteins

Protein lysates were loaded onto Pierce™ High-Capacity Streptavidin Agarose (Thermo Fisher Scientific, Waltham, Massachusetts, USA). 100 μL and 150 μL packed bead volumes were used for epimastigote and amastigote samples, respectively, and incubated on a rotative wheel overnight at 4°C. For all following steps, washes consisted of adding 1 mL of the indicated buffer and placing the tube on the rotative wheel for 5 minutes, then spinning down the agarose beads for 1 minute at 500 x g, as previously described Beads with bound protein were subjected to 5 washes with Buffer 1 (8 M urea, 200 mM NaCl, 100 mM Tris, pH 8.0) with 0.2% sodium dodecyl sulfate (SDS), 5 washes with Buffer 1 containing 2% SDS, and 5 washes with Buffer 1 with no SDS were completed at room temperature. Next, 2 washes with 200 mM NaCl, 100 mM Tris, at a pH of 7.0 and 2 washes with Tris, pH 8.0 were completed at 4°C. Washed beads were adjusted to pH 7.5 with 200 mM HEPES (4-(2-hydroxyethyl)-1-piperazineethanesulfonic acid) and bound proteins were reduced using 5 mM dithiothreitol (Sigma-Aldrich) at 37°C for 1 h, followed by alkylation of cysteine residues using 15 mM iodoacetamide (Sigma-Aldrich) in the dark at room temperature for 1 h. Excessive iodoacetamide was quenched using 10 mM dithiotheritol. Protein mixtures were diluted in 1:6 ratio (v/v) using ultrapure water prior to digestion using sequencing grade trypsin (Promega) at 37°C for 16 h. Digested peptides were subsequently desalted using self-packed C18 STAGE tips (3M EmporeTM) ^57^ for LC-MS/MS analysis.

### Mass spectrometry

#### Epimastigotes

Desalted peptides were resuspended in 0.1% (v/v) formic acid and loaded onto HPLC-MS/MS system for analysis on an Orbitrap Q-Exactive Exploris 480 (Thermo Fisher Scientific) mass spectrometer coupled to an Easy nanoLC 1000 (Thermo Fisher Scientific) with a flow rate of 250 nl/min. The stationary phase buffer was 0.5 % formic acid and mobile phase buffer was 0.5 % (v/v) formic acid in acetonitrile. Chromatography for peptide separation was performed using increasing organic proportion of acetonitrile (5 – 40 % (v/v)) over a 120 min gradient) on a self-packed analytical column using PicoTipTM emitter (New Objective, Woburn, MA) using Reprosil Gold 120 C-18, 1.9 μm particle size resin (Dr. Maisch, Ammerbuch-Entringen, Germany). The mass spectrometry analyzer operated in data dependent acquisition mode with a top ten method at a mass range of 300–2000 Da. Data were processed using MaxQuant software (version 1.5.2.8) ^58^ with the following setting: oxidized methionine residues and protein N-terminal acetylation as variable modification, cysteine carbamidomethylation as fixed modification, first search peptide tolerance 20 ppm, main search peptide tolerance 4.5 ppm. Protease specificity was set to trypsin with up to 2 missed cleavages allowed. Only peptides longer than five amino acids were analyzed, and the minimal ratio count to quantify a protein is 2 (proteome only). The false discovery rate (FDR) was set to 1% for peptide and protein identifications. Database searches were performed using the Andromeda search engine integrated into the MaxQuant environment ^59^ against the UniProt *Trypanosoma cruzi* strain CL Brener (352153) database containing 19,242 entries (March 2020). “Matching between runs” algorithm with a time window of 0.7 min was employed to transfer identifications between samples processed using the same nanospray conditions. Protein tables were filtered to eliminate identifications from the reverse database and common contaminants.

#### Amastigotes

Desalted peptides were resolubilized in 0.1% (v/v) formic acid and loaded onto HPLC-MS/MS system for analysis on an Orbitrap Q-Exactive Exploris 480 (Thermo Fisher Scientific) mass spectrometer coupled to an FAIMS Pro Interface system and Easy nanoLC 1000 (Thermo Fisher Scientific) with a flow rate of 300 nl/min. The stationary phase buffer was 0.1 % formic acid, and mobile phase buffer was 0.1 % (v/v) formic acid in 80% (v/v) acetonitrile. Chromatography for peptide separation was performed using increasing organic proportion of acetonitrile (5 – 40 % (v/v)) over a 120 min gradient) on a self-packed analytical column using PicoTip™ emitter (New Objective, Woburn, MA) using Reprosil Gold 120 C-18, 1.9 μm particle size resin (Dr. Maisch, Ammerbuch-Entringen, Germany). High precision iRT calibration was used for samples processed using the same nanospray conditions ^60^. The mass spectrometry analyzer operated in data independent acquisition mode at a mass range of 300–2000 Da, compensation voltages of −50/-70 CVs with survey scan of 120,000 and 15,000 resolutions at MS1 and MS2 levels, respectively. Data were processed using Spectronaut™ software (version 15; Biognosys AG) ^61^ using directDIA™ analysis with default settings, including: oxidized methionine residues, biotinylation, protein N-terminal acetylation as variable modification, cysteine carbamidomethylation as fixed modification, initial mass tolerance of MS1 and MS2 of 15 ppm. Protease specificity was set to trypsin with up to 2 missed cleavages were allowed. Only peptides longer than seven amino acids were analyzed, and the minimal ratio count to quantify a protein is 2 (proteome only). The false discovery rate (FDR) was set to 1% for peptide and protein identifications. Database searches were performed against the UniProt *Trypanosoma cruzi* strain CL Brener (352153) database containing 19,242 entries (March 2020). Protein tables were filtered to eliminate identifications from the reverse database and common contaminants.

### Principal component analysis

All protein intensity scores were uploaded to Metaboanalyst 5.0 ^62^ to perform statistical analysis, with one factor. Data was entered as peak intensities and filtered using the interquartile range, then normalized by sum and log transformed. 2D PCA scores were plotted in Prism GraphPad.

### Volcano plots

For epimastigote data, protein intensity scores for all proteins that were found to be 100-fold or higher enriched in the F-Turbo or C-Turbo samples over wild type were loaded into Prism GraphPad, and Log_10_ transformed. Multiple unpaired t-tests were run on the data with a false discovery rate of 1% and the results were reported as F-Turbo – C-Turbo, with -log10(q value) reported for the volcano plot. For amastigote data, Spectronaut™ software was used to generate the statistical analysis of the amastigote proteomics. Within the DIA analysis pipeline, the default settings were used, including a false discovery rate of 1% and a n unpaired Student’s t-test was performed. Fold changes were reported as an average Log10 (epimastigotes) or Log2 (amastigotes) ratio for the volcano plot. Volcano plot was created in Prism GraphPad. Post-analysis, six proteins were excluded from the amastigote flagellar enriched list, as they were not identified in 2 of 3 biological replicates. Additionally, all allelic duplicates were removed to ensure that the proteins listed in the final tables were represented by a single gene identifier corresponding to the CL Brener reference genome (TriTrypDB; https://tritrypdb.org/), but paralogs remain.

### Interpro domain enrichment analysis

DAVID ^63^ enrichment was used to assign Interpro domains to all of the proteins found to be significantly enriched in either of the F-Turbo samples. No weighting of the data was completed. Interpro domains with a p-value ≤ 0.05 were considered significantly enriched.

### Indirect immunofluorescence microscopy

Epimastigotes were fixed directly in growth medium with the addition of paraformaldehyde (1% final concentration in PBS) for 10-minute at 4°C. Fixed parasites were pelleted by centrifuging for 10 minutes at 4000 x g and resuspended in PBS. 10 μL of the parasite solution was dropped onto poly-L-lysine coated slides and allowed to dry completely prior to staining. For immunostaining of intracellular amastigotes, *T. cruzi*-infected NHDF on round cover glass (12 mm, #1.5; Electron Microscopy Sciences, Hatfield, Pennsylvania, USA) were fixed at 48 hours post-infection with 1% (v/v) paraformaldehyde/PBS. All steps of the immunostaining protocol were preceded by three washes with PBS and carried out at room temperature. Parasites were permeabilized with a 0.1% Triton-X 100 solution (JT Baker, Phillipsburg, New Jersey, USA) for 10 minutes and blocked with 3% BSA (Sigma-Aldrich, St. Louis, Missouri, USA) in PBS for 1 hour. The primary antibody solution containing 1:400 mouse α-FLAG (clone M2, Sigma-Aldrich, St. Louis, Missouri, USA) and/or 1:1,500 rabbit α-FCaBP ^64^ in 1% BSA in PBS was added for 1 hour, followed by a 1:1000 α-Mouse Alexa Flour 594 and/or α-Rabbit Alexa Flour 647 solution in 1% BSA in PBS for 1 hour. DAPI (0.2 μg/mL; Thermo Fisher Scientific, Waltham, Massachusetts, USA) in PBS was added for 5 minutes, and following washes, coverslips were placed onto slides with Prolong® Diamond mountant (Thermo Fisher Scientific, Waltham, Massachusetts, USA). After setting for 24 hours parasites were imaged using a Yokogawa CSU-X1 spinning disk confocal system paired with a Nikon Ti-E inverted microscope and an iXon Ultra 888 EMCCD camera (100X objective). Image processing, analysis, and display were performed using ImageJ Fiji software^65^.

## Acknowledgments

We thank the Sabri Ülker Center’s Advanced Imaging Lab at the Harvard T.H. Chan School of Public Health for microscopy training and access. Additionally, we would like to thank Zon Weng Lai from the Harvard Chan Advanced Multi-omics Platform at the Harvard T.H. Chan School of Public Health for his assistance in interpreting the mass spectrometry data. We thank Jeffrey Dvorin (Boston Children’s Hospital) for providing the TurboID plasmid, David Engman (Cedars-Sinai Medical Center) for the gift of the anti-FCaBP antibody and Jay Bangs (The State University of New York at Buffalo) for anti-BiP antibody. We also thank Lucas Pagura for his thoughtful discussion. This work was supported by NIH grant R21AI135520 awarded to BAB and NIH training grant 5T32AI049928.

---

**Supplementary Figure 1.**
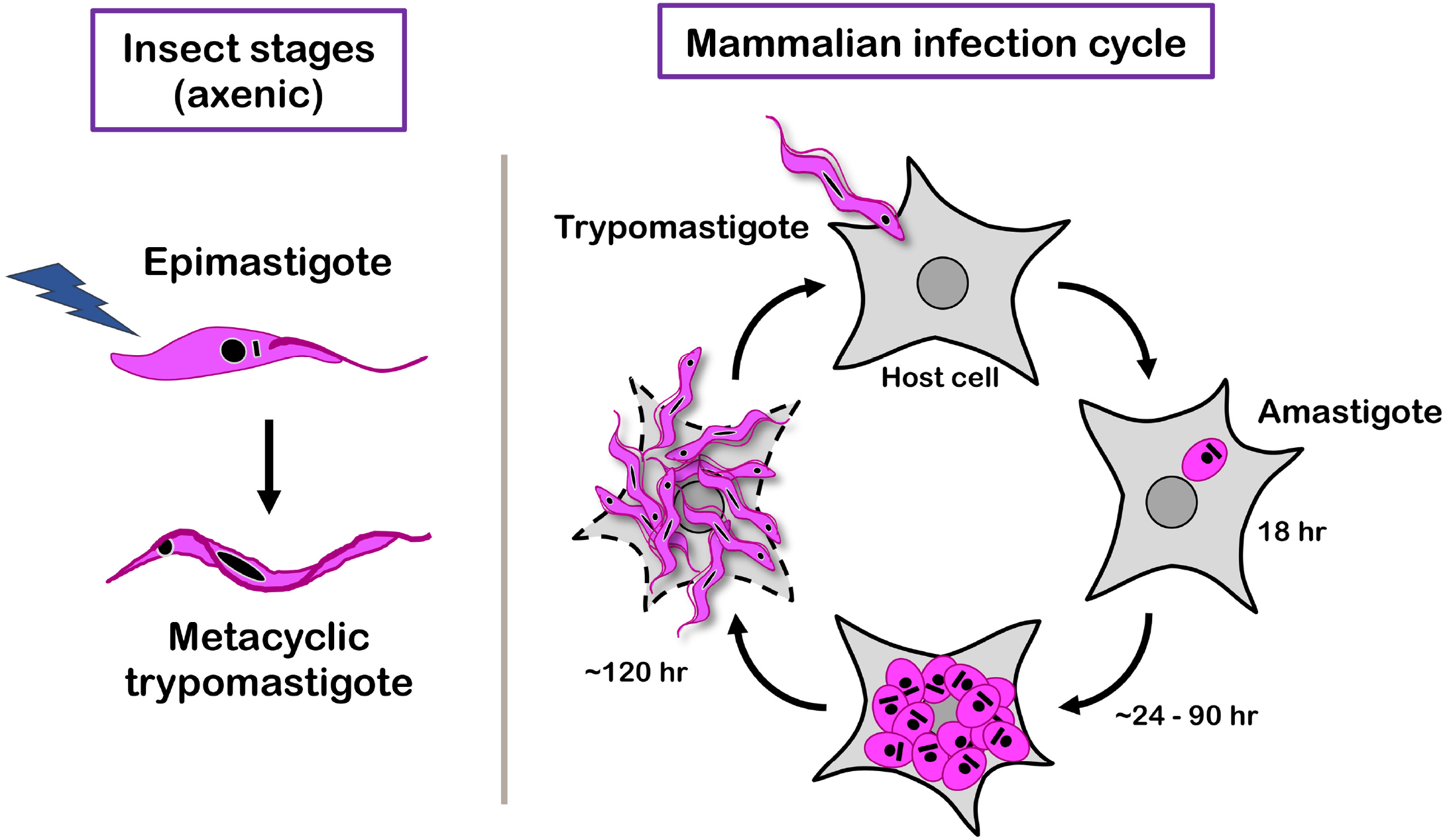
Model of the *T. cruzi* life cycle. Schematic of the *T. cruzi* life cycle highlighting the insect stage *‘epimastigote’* that is propagated axenically in liquid culture and gives rise to the infectious *‘metacyclic trypomastigote’*. Trypomastigotes, whether derived from epimastigotes or as the end product of a single lytic cycle in a mammalian cell are motile, non-dividing forms of the parasite that actively invade a mammalian host cell. Inside a host cell, the *‘trypomastigote’* transforms into the replicative intracellular *‘amastigote’* stage by 18 hours post-infection (hpi). Amastigotes undergo several rounds of proliferation, dividing by binary fission (between ~24-90 hpi), before they stop dividing and differentiate into trypomastigotes, that eventually lyse the infected host cell and disseminate infection. Stable transfection and drug selection is performed in the epimastigote stage (lightning bolt symbolizes electroporation). Once stable genomic changes are confirmed in epimastigotes, these parasites are used to establish the mammalian infection cycle starting with metacyclic trypomastigotes as outlined above.

**Supplementary Figure 2:**
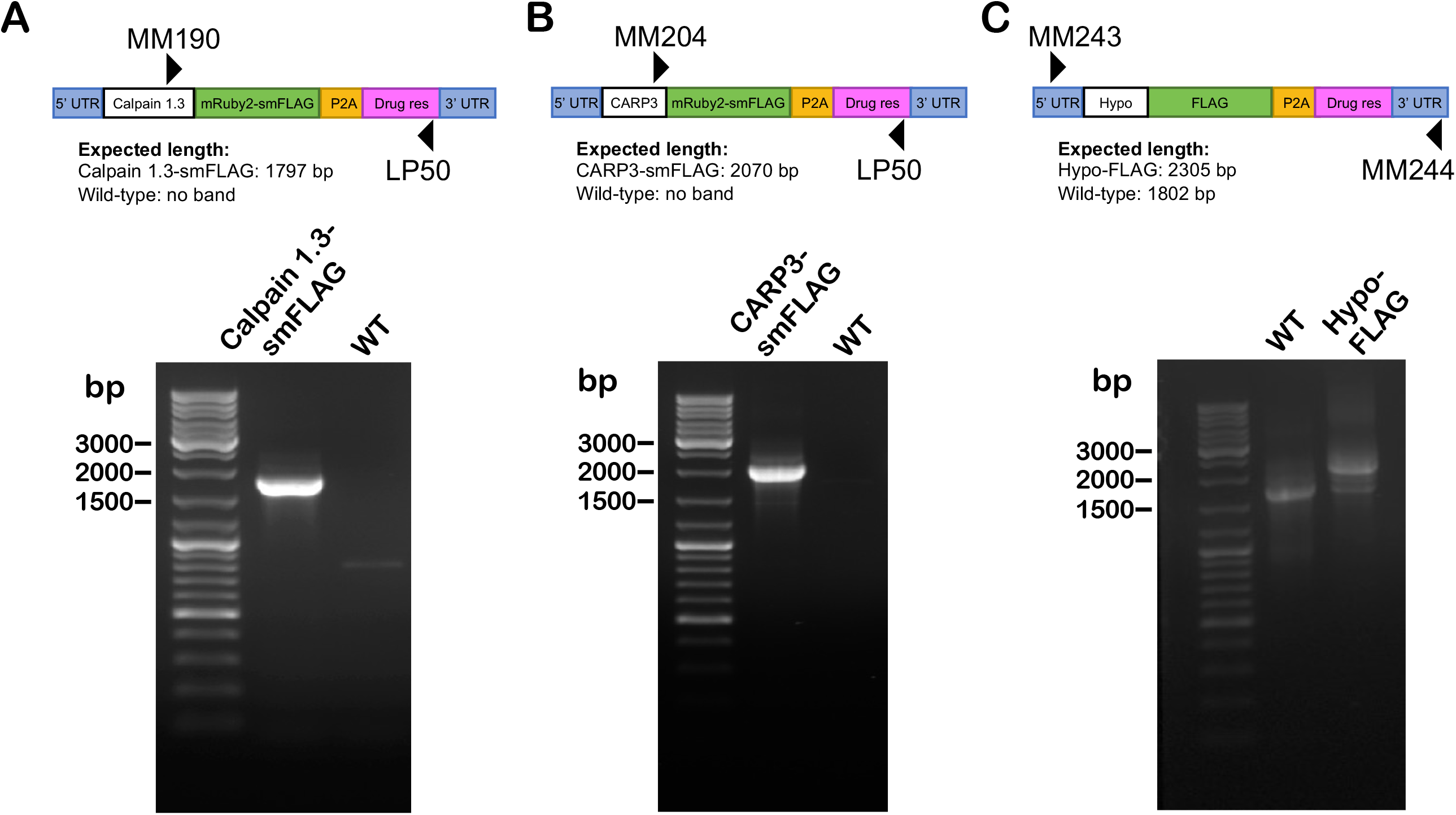
PCR confirmation of endogenous tags for candidate proteins. **A.** Schematic showing the region of amplification and DNA gel with corresponding bands for **A**. calpain 1.3-smFLAG (TcCLB.506563.200), **B.** CARP3-smFLAG (TcCLB.506681.40), and **C.** hypothetical protein-FLAG (TcCLB.510329.180). Ladder run on all DNA gels is Thermo Scientific GeneRuler DNA Ladder.

**Supplementary Table 1: Raw proteomics data for epimastigotes and intracellular amastigotes TurboID experiment.** Each sheet contains the protein identity and intensity score for all samples in either epimastigotes or amastigotes.

**Supplementary Table 2: Filtered proteomics data for epimastigotes and intracellular amastigotes TurboID experiment.** Each sheet contains the protein identity and intensity score for all samples in either epimastigotes or amastigotes that was used for statistical analysis.

**Supplementary Table 3: Proteins enriched in the flagellar TurboID expressing samples for epimastigotes and intracellular amastigotes**. Each sheet contains information about the proteins enriched in either epimastigotes, amastigotes, or both epimastigotes and amastigotes.

**Supplementary Table 4: Primers for all PCR and endogenous tagging**. Primer pairs are listed for all experiments described.

